# Hibernation shows no apparent effect on germline mutation rates in grizzly bears

**DOI:** 10.1101/2022.03.15.481369

**Authors:** Richard J. Wang, Yadira Peña-Garcia, Madeleine Bibby, Muthuswamy Raveendran, R. Alan Harris, Heiko T. Jansen, Charles T. Robbins, Jeffrey Rogers, Joanna L. Kelley, Matthew W. Hahn

## Abstract

A male mutation bias is observed across vertebrates, and, where data are available, this bias is accompanied by increased per-generation mutation rates with parental age. While continuing mitotic cell division in the male germline post-puberty has been proposed as the major cellular mechanism underlying both patterns, little direct evidence for this role has been found. Understanding the evolution of the per-generation mutation rate among species requires that we identify the molecular mechanisms that change between species. Here, we study the per-generation mutation rate in an extended pedigree of the brown (grizzly) bear, *Ursus arctos horribilis*. Brown bears hibernate for one-third of the year, a period during which spermatogenesis slows or stops altogether. The cessation of spermatogenesis is predicted to lessen the male mutation bias and to lower the per-generation mutation rate in this species. However, using whole-genome sequencing, we find that both male bias and per-generation mutation rates are the same as expected for a non-hibernating species. We also carry out a phylogenetic comparison of substitution rates along the lineage leading to brown bear and panda (a non-hibernating species) and find no slowing of the substitution rate in the hibernator. Our results contribute to accumulating evidence that suggests that male germline cell division is not the major determinant of mutation rates and mutation biases. The results also provide a quantitative basis for improved estimates of the timing of carnivore evolution.

## Introduction

The per-generation mutation rate evolves between species. Whole-genome sequencing has revealed that the mutation rate varies by several orders of magnitude across eukaryotes (Lynch 2010) and by at least two-fold among mammals (Chintalapati and Moorjani 2020; Wang et al. 2021a). Prior to the advent of large-scale DNA sequencing, early studies of disease mutations in humans uncovered two general patterns in the accumulation of mutations. First, parental age was found to be positively correlated with the probability of observing a mutation: older parents were more likely to have children with inherited diseases (Weinberg 1912; Risch et al. 1987). Second, advanced paternal age better predicted the appearance of disease mutations than advanced maternal age—implying that mutation was male-biased (Haldane 1946; Penrose 1955). Both of these patterns have been confirmed by studies that have sequenced large numbers of human pedigrees (Kong et al. 2012; Rahbari et al. 2016; Goldmann et al. 2016; Jónsson et al. 2017), as well as by studies in other mammals (Venn et al. 2014; Thomas et al. 2018; Besenbacher et al. 2019; Lindsay et al. 2019; Wang et al. 2020; Wu et al. 2020; Bergeron et al. 2021; Wang et al. 2021a).

The main mechanism proposed to explain both the parental age effect and male-biased mutation is the continuing replication of the male germline in mammals. After a relatively fixed number of mitotic divisions during germline development before puberty in both sexes, the male germline continues to undergo mitotic cell division post-puberty (Drost and Lee 1995). Although the polymerases responsible for genome replication have very low error rates (McElhinny et al. 2010a,b), the cell lineage leading to any given spermatozoon can go through hundreds of spermatogenic cell divisions. This difference in the contribution of mutations between males and females informs evolutionary models of the mutation rate, which usually combine a period of constant pre-puberty mutation accumulation in both sexes with a period of increasing mutation post-puberty only in males (e.g., Thomas and Hahn 2014; Amster and Sella 2016; Gao et al. 2016; Thomas et al. 2018). While there is a small effect of environmental damage on the accumulation of mutations in both males and females with age (e.g., Goldmann et al. 2016; Jónsson et al. 2017), very large samples sizes have been needed to detect it; therefore, it is often ignored in these models.

Despite the general acceptance of the male germline replication model of mutation accumulation, several patterns from whole-genome sequencing studies have emerged that do not fit easily within this paradigm. Here we mention four of these patterns (see de Manuel et al. 2022 for further discussion). 1) Spermatogenic cycle length is not predictive of mutation rates: non-human primates with shorter seminiferous epithelial cycle lengths do not show a faster rate of mutation accumulation per year (Wang et al. 2020; Wu et al. 2020). Although the number of these cell cycles may not exactly match the number of replications (Scally 2016; Thomas et al. 2018), shorter cycle lengths should result in more mutations per unit time. 2) In humans, the male bias in mutations exists even in the youngest fathers studied (Gao et al. 2019). If male bias is largely driven by the continued replications of spermatogenesis post-puberty, there should be little difference between the sexes immediately post-puberty. 3) CàT mutations at CpG sites show a similar degree of male bias as other mutations (Jónsson et al. 2017), even though they are not thought to be associated with polymerase errors during replication. While both sexes can incur exogenous damage at these sites, it is not clear how or why such damage would cause more mutations in males. 4) Studies of somatic mutagenesis have not found a higher mutation rate in tissues that are mitotically active (Abascal et al. 2021). While variation in the somatic mutation rate across tissue types was observed, the mutation rate was not associated with the rate of cellular division in each tissue.

With the growing number of questions surrounding the role of male germline replication in the evolution of the mutation rate, species that experience a slowdown or cessation of this replication represent a potentially illuminating study system. Many mammals (and non-mammals) undergo a period of quiescence or torpor during the winter—what is generally referred to as hibernation (Geiser 2013). Many physiological functions are altered during hibernation; in particular, reproduction and the activity of reproductive tissues are minimal during hibernation, with a complete cessation of spermatogenesis in some species (e.g., in black bears [*Ursus americanus*]; Tsubota et al. 1997). Male germline activity restarts in late winter through a process known as testicular recrudescence.

The brown (grizzly) bear, *Ursus arctos horribilis*, is a model system for studies of the genetics and physiology of mammalian hibernation (Hershey et al. 2008; McGee et al. 2008; Buffenstein et al. 2014; Rigano et al. 2017; Jansen et al. 2019; Mugahid et al. 2020). During hibernation, bears do not eat, produce minimal-to-no urine, reduce heart rates to 10-15 beats per minute, do not lose bone mass, and have minimal muscle mass loss despite having almost no weight-bearing activity. Brown bears are seasonal breeders, with the peak breeding season occurring in June. After breeding season, the testis becomes reduced in size, and—at least in closely related black bears— spermatogenesis and reproductive steroidogenesis are greatly reduced as the male enters hibernation (Howell-Skalla et al. 2000). Our attempts to obtain sperm from hibernating brown bears via electrostimulation have been unsuccessful, further supporting the idea that spermatogenesis is suspended during hibernation.

Given the annual pause in male germline replication experienced by brown bears, we hypothesized that the per-generation mutation rate and the degree of male bias in mutations from this species would be lower under the male germline replication model of mutation accumulation. Here, we test this hypothesis by studying the mutation rate in an extended pedigree of brown bears. Using whole-genome sequencing of the four trios embedded in this pedigree, we find no difference between our estimate of the per-generation mutation rate and its expectation under a model without hibernation. We also find no difference in the degree to which mutations are male-biased compared to other mammals. Further analysis of the per-year mutation rate—estimated via phylogenetic comparison with closely related non-hibernating species—also shows no effect of hibernation. We discuss the implications of these results for our understanding of the cellular basis of mutation rate evolution in mammals.

## Methods

### Animals

Brown bears (*Ursus arctos horribilis* Linnaeus 1758) were housed at the Washington State University Bear Research, Education and Conservation Center (WSU Bear Center, Pullman, WA, USA) in accordance with the Bear Care and Colony Health Standard Operating Procedures approved by the Washington State Institutional Animal Care and Use Committee (IACUC) protocol #6546. The bears at WSU Bear Center hibernate from November to mid-to-late March.

### Testes measurements

Two adult males were periodically anesthetized as previously described (Ware et al. 2012) over a four-year period. Both males were 13 years old at the time of first measurement. The final dataset includes measurements at roughly monthly intervals between January and December, though each bear was only measured approximately three times in any particular year. Once each bear was anesthetized, each testis was manually palpated and externalized with gentle pressure. Paired testes measurements, including skin, were then made using a caliper micrometer (Mitutoyo, model 505-681) to the nearest 0.1 mm. The length (L) and width (W) of the testes were measured three times, and the average values for each testis were recorded. An estimated testis volume was then derived for each testis using the formula, W^2^ × L (as described by Gorman and Zucker 1995), and the two testis values were added together to generate a total estimated testis volume per individual.

### DNA Extraction and Quantification

Samples from an extended pedigree with four embedded trios (*n*=8 individuals; Figure 1) were used for per-generation mutation rate estimates. Blood was collected (~5 ml) from the jugular vein into PAXgene Blood DNA Tubes. DNA was extracted using the PAXgene Blood DNA Kit following the standard protocol for whole blood with no modifications. DNA was quantified with the high sensitivity double stranded (ds) DNA Assay Kit (Qubit dsDNA HS Assay Kit, Thermo Fisher Scientific, #Q32854) on the Qubit 2.0 fluorometer.

**Figure 1.**
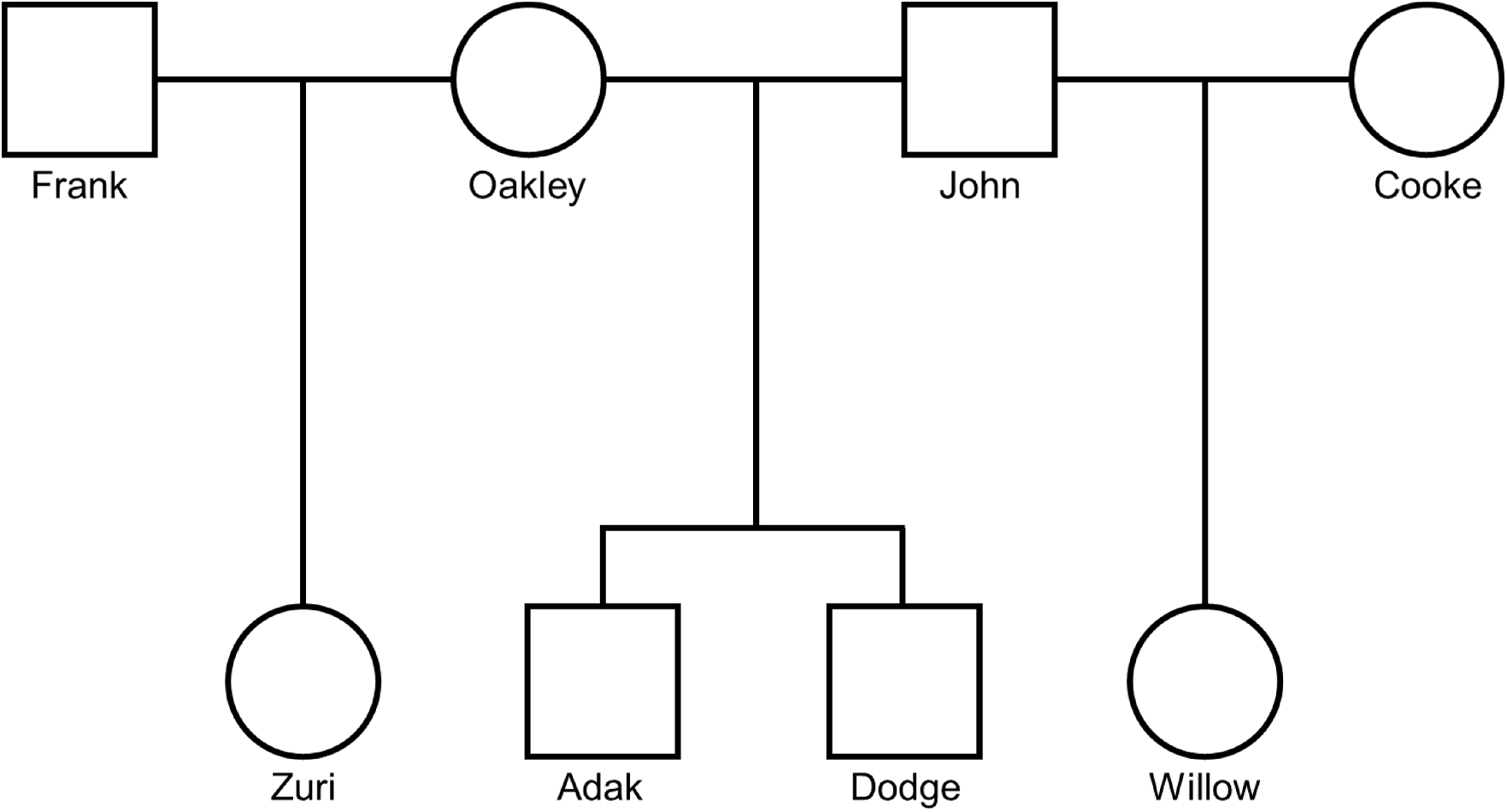
Pedigree of bears included in the study. Eight individuals that were part of an extended pedigree were sequenced. The four probands (Zuri, Adak, Dodge, and Willow) each represents the offspring within an independent trio. Males are indicated by squares and females by circles.

### DNA Sequencing

Extracted DNA was sequenced at the Baylor College of Medicine Human Genome Sequencing Center (Houston, Texas). Standard PCR-free libraries were prepared using KAPA Hyper PCR-free library reagents (KK8505, KAPA Biosystems). Total genomic DNA was sheared into fragments of approximately 200-600 base pairs (bp) and purified using AMPure XP beads. Sheared DNA molecules were subjected to double size-selection with different ratios of AMPure XP beads to select a narrow size band of sheared DNA molecules for library preparation. This was followed by DNA end-repair and 3’-adenylation before the ligation of barcoded adapters. Library quality was evaluated by fragment analysis and qPCR assay. The resulting libraries were sequenced on an Illumina NovaSeq 6000, producing 150 bp paired-end reads.

### Mutation identification

Sequenced reads were aligned with BWA-MEM version 0.7.12-r1039 (Li 2013) to the domestic brown bear reference genome, ASM358476v1 (Taylor et al. 2018). Picard MarkDuplicates v. 1.105 (Broad Institute 2019) was used to identify and mark duplicate reads from the BAM files. We used GATK v. 4.1.2.0 (Van der Auwera et al. 2013) to call variants using best practices. HaplotypeCaller was used to generate gVCF files for each sample and joint genotype calling across samples was performed with GenotypeGVCFs. We applied GATK hard filters: (SNPs: “QD < 2.0 || FS > 60.0 || MQ < 40.0 || MQRankSum < −12.5 || ReadPosRankSum < −8.0”) and removed calls that failed.

We used the same pipeline for identifying autosomal *de novo* mutations from called variants as in our previous work (Wang et al. 2020; Wang et al. 2021a), which we summarize here: An initial set of candidate mutations was identified as “Mendelian violations” in each trio. Specifically, we looked for violations where both parents were reference homozygous and the offspring was heterozygous for an alternate allele. As this is the most common type of genotyping error (Wang et al. 2021b), we then apply the following filters to the initial set of candidates to get a set of high-confidence candidates:

1. Read-depth at the candidate site must be between 20 and 80 for every individual in the trio. Sites with too few reads are likely to be sampling errors, while sites with too many reads are likely to be from repetitive regions.
2. High genotype quality (GQ) in all individuals (GQ > 60).
3. Candidate mutations must be present on reads from both the forward and reverse strand in the offspring.
4. Candidate mutations must not be present in any reads from either parent.
5. Candidate mutations must not be present in any other samples (except siblings).
6. Candidate mutation must not have low allelic depth in the offspring (allelic balance > 0.30).

We assessed the sensitivity of our mutation rate estimates across a range of stringency criteria and found them to be in good agreement across reasonable filter limits (Figure S1). The distribution of allelic balances was also centered at 0.5 (Figure S2).

### Per-generation mutation rate estimate

In order to transform the identified number of *de novo* mutations into a rate per-base per-generation, we need an accurate count of the number of bases at which mutations could have been identified in each trio. As in previous work, we applied existing strategies that considered differences in coverage and filtering among sites (Besenbacher et al. 2019; Wang et al. 2020; Wang et al. 2021a), and that estimate false negative rates from said filtering. Briefly, the number of identified mutations was divided by the total number of “callable sites.” Callable sites are a product of the number of sites covered by the appropriate sequencing depth and the estimated probability that such a site would be called correctly given that it was a true *de novo* mutation. The mutation rate is then calculated as:

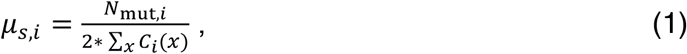

where *μ_s,i_* is the per-base mutation rate for trio *i*, *N*_mut,*i*_ is the number of mutated bases in trio *i*, and *C_i_*(*x*) is the callability of site *x* in that trio. This strategy assumes that the ability to call each individual in the trio correctly is independent, allowing us to estimate *C_i_* (*x*) as:

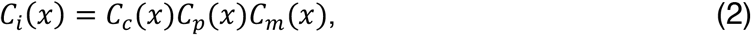

where *C_c_*, *C_p_*, and *C_m_* are the probability of calling the child, father, and mother correctly for trio *i*. These values are estimated by applying the same set of stringent filters to high-confidence calls from each trio. For heterozygous variants in the child,

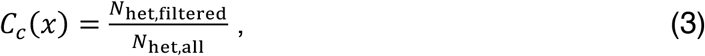

where *N*_het,all_ is the number of variants in the offspring where one parent is homozygous reference and the other parent is homozygous alternate, leading to high confidence in the child heterozygote call, and *N*_het,filtered_ is the set of such calls that pass our child-specific candidate mutation filters. The parental callability, *C_p_*(*x*) and *C_m_*(*x*), were estimated in a similar manner, by calculating the proportion of remaining sites in each after the application of the stringent mutation filters. Based on our previous results and the results of comparisons of our pipeline to those from other research groups when applied to common datasets (Bergeron et al. 2022), we assume our pipeline produces no (or very few) false positives.

### Phasing mutations

We used read-pair tracing to determine the parent of origin (“phase”) for mutations across all of our trios. We did this by applying WhatsHap 1.0 (Patterson et al. 2015) in read-based phasing mode for each individual separately, and then matched informative blocks bearing the mutation to their parent of origin according to the rules of Mendelian inheritance. Ambiguous blocks, including any that showed genotype inconsistencies between parent and offspring, were left unphased.

### Per-year mutation rate estimate

To estimate long-term rates of molecular evolution, we identified orthologs from brown bear (ASM358476v1), panda (*Ailuropoda melanoleuca*, ASM200744v2), dog (*Canis lupus familiaris*, Cfam_1.0), and ferret (*Mustela furo*, MusPutFur1.0) using OrthoFinder v. 2.5.2 (Emms and Kelly 2019) with DIAMOND v. 0.9.27 (Buchfink et al. 2021) as the sequence search program. Only orthogroups with single-copy orthologs were considered in the analysis. We were not able to confidently place genes on the bear X chromosome, but excluded the set of genes with human orthologs on the X from all further comparisons.

Protein-coding sequences for each orthogroup containing all four species were aligned by codon using GUIDANCE2 (Sela et al. 2015) with MAFFT v. 7.471 (Katoh and Standley 2013). GUIDANCE2 provides quality scores for each residue and column of the alignment. Scores were used to remove unreliable sequence: low-confidence residues with scores <0.93 were converted into gaps. Columns with gaps and N’s were removed from the alignments using trimAl v. 1.4.rev22 (Capella-Gutierrez et al. 2009). Additionally, alignments with sequences that were shorter than 200 bp were filtered. This process resulted in a total of 4,886 gene alignments that were considered for further analysis.

Synonymous substitutions per site (*d*s) for each branch were estimated using HyPhy (Pond et al. 2005) with the FitMG94.bf model (https://github.com/veg/hyphy-analyses). We assumed that every gene had the same topology: (((*U. a. horribilis, A. melanoleuca), M. furo), C. l. familiaris*).). The average tip branch lengths leading to brown bear and panda were obtained by taking the mean of *d*s values across all genes, after removing genes where either of the two tip branches were longer than 0.2 (which we took as an indication of poor alignment). Despite the absence of an absolute time estimate for the split between brown bear and panda, their comparison as sister lineages provides an equal amount of time for substitutions to have accumulated in each species, and we therefore refer to the estimated distances as substitution rates.

### Expected per-generation mutation rate in the absence of hibernation

To compare the estimated per-generation mutation rate obtained in our bear pedigrees to that expected in a non-hibernating species that is otherwise equivalent, we used the reproductive longevity model described in Thomas et al. (2018). This model divides the per-generation mutation rate into the contributed rate from each of three different life stages: (1) female (*μ_gF_*), (2) male before puberty (*μ_gM0_*), (3) and male after puberty (*μ_gM1_*). The first two parameters are constants, while the third is a function of paternal age post-puberty. Our goal is to estimate the total per-generation mutation rate (*μ_g_*) as a function of these three parameters. Since autosomes spend half their time in males and half their time in females, this relationship becomes:

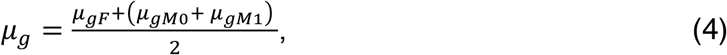

where the two values coming from males are grouped together in the numerator (equation 8 in Thomas et al. 2018). We estimate *μ_gM1_* as a function of the yearly rate of mutation accumulation in males post-puberty (*μ_yM1_*) and the reproductive longevity (*RL*) of the father:

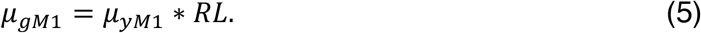

Specifically, we calculate *RL* as the difference between the age of puberty in males (*P_M_*) and the age of the male parent at conception of his offspring (*A_M_*):

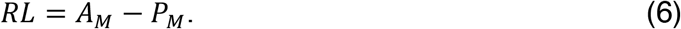

*RL* therefore accounts for the amount of time mutations have had to accumulate postpuberty in males, while *μ_gM1_* describes the number of such post-puberty mutations.

Values for the age at puberty and conception are used for the bear species considered (see Results), and we use estimated values of *μ_gF_, μ_gM0_*, and *μ_yM1_* from previous work: 14.2 mutations, 15.5 mutations, and 2.01 mutations/year, respectively (Kong et al. 2012; Thomas et al. 2018). Although these calculations assume that many parameter values are the same between humans and bears, we have found them to be remarkably well-conserved across species (Thomas et al. 2018; Wang et al. 2020; Wang et al. 2021a).

### Expected per-year mutation rate in the absence of hibernation

The model described above can be extended to calculate the expected per-*year* mutation rate as a function of differing life histories (Thomas and Hahn 2014; Segurel et al. 2014; Amster and Sella 2016; Gao et al. 2016). We used the above estimates along with parameters from the life histories of brown bears and pandas to calculate the expected per-year mutation rate, without regard for hibernation status.

To calculate mutation rates per year (*μ_y_*) we sum the mutational contribution from each life stage per generation, and weight these contributions by the amount of time spent in each:

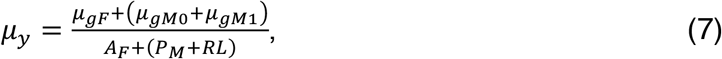

where *A_F_* is the age of the female parent at conception, and all other parameters are the same as defined above (equation 9 in Thomas et al. 2018). Although some terms could be simplified, writing the equation out in this way helps to clarify which mutational parameters correspond to which sex and to which developmental stage.

## Results

### Testes size through hibernation

To highlight the physiological and phenotypic changes that the male germline undergoes during hibernation, we measured seasonal variation in testis size from two sexually mature male brown bears across a four-year period (Figure 2). The results show clear seasonal differences with a testicular recrudescence (regrowth) evident during hibernation and reduction in size (regression) during the hyperphagia period (August – October). These results confirm other observations of seasonal changes in testis size and function made in both brown and black bears (Tsubota and Kanagawa 1989; Hellgren 1998; White et al. 2005; Spady et al. 2007). Moreover, our electrostimulation attempts during hibernation were unsuccessful, further supporting the idea that spermatogenesis is dramatically reduced or suspended during hibernation.

**Figure 2.**
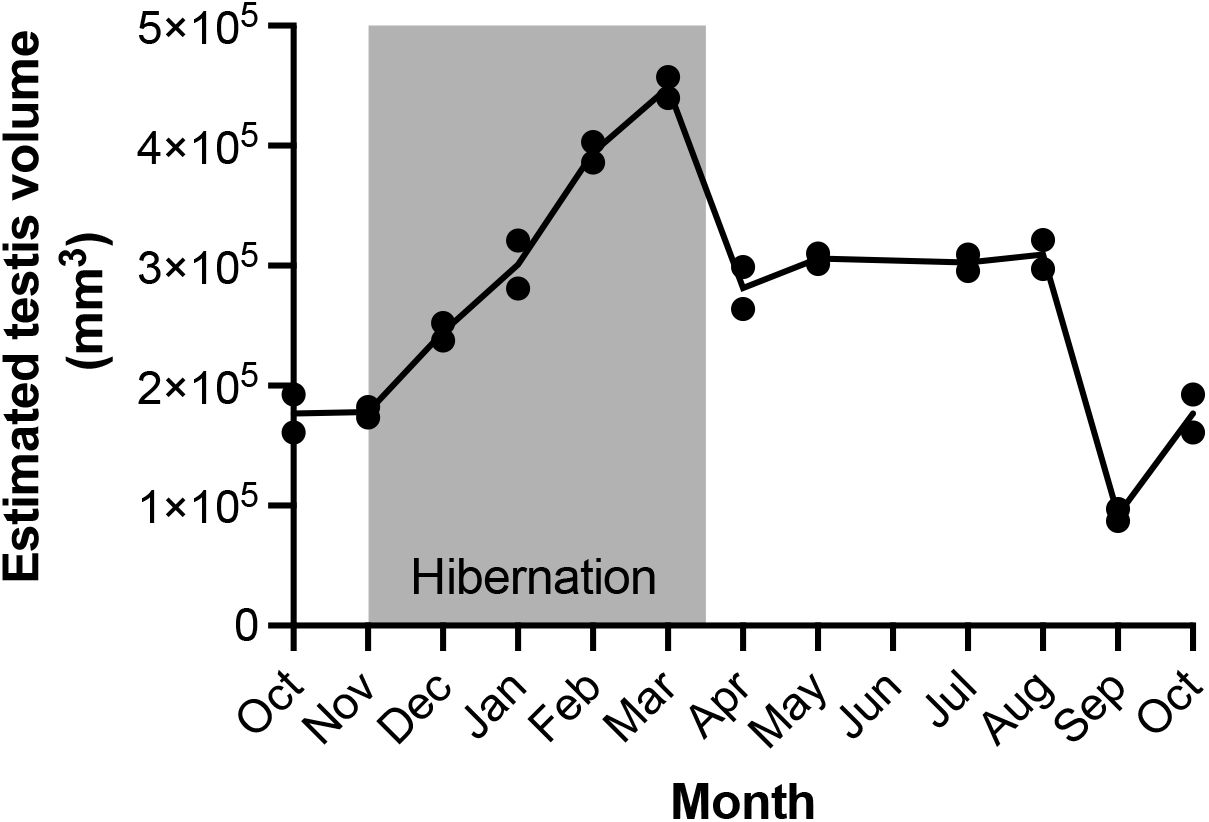
Estimated testis volume through hibernation. Two male grizzly bears were sampled so that each bear was measured in each month of the year at least once (measurements were spread across a four-year period). Dots indicate individual values, and the line is the mean volume. Grey shading indicates the timing of hibernation.

### Estimating the per-generation mutation rate

We sequenced eight individuals from a large pedigree of captive brown bears kept at the Washington State University Bear Center (Figure 1). Individual samples had an average of 51.1 × coverage (min: 46.6×, max: 57.1 ×), with reads mapped to the brown bear reference genome (NCBI assembly ASM358476v1). The pedigree can be separated into four trios from which independent mutation rate estimates can be made (we observed no candidate mutations shared among siblings). We required that all three individuals in a trio have a minimum (and maximum) depth of high-quality reads for a mutation to be called (Methods). On average, these filters for “callability” allowed us to examine 1.72 Gb per trio for mutation identification (Table 1).

**Table 1.**

| Trio | Proband | Mutations | Callable size (Mb) | Per bp rate ($\times 10^{-8}$) | Parental age (y) | |
| --- | --- | --- | --- | --- | --- | --- |
|  |  |  |  |  | Paternal | Maternal |
| 1 | Zuri | 31 | 1753 | 0.88 | 13 | 12 |
| 2 | Adak | 23 | 1701 | 0.68 | 13 | 12 |
| 3 | Dodge | 31 | 1678 | 0.92 | 13 | 12 |
| 4 | Willow | 30 | 1752 | 0.89 | 13 | 12 |

Mean mutation rate: $0.84 \times 10^{-8}$ per bp per generation

After applying a stringent set of filters, we identified 115 total mutations across the four trios, including one multinucleotide mutation (Supplementary Table 1). All of the trios have parents that are the same ages (to the nearest year) and consequently we found very little variation in the number of mutations per trio (Table 1). To estimate the per base pair mutation rate, we divided the number of mutated bases identified in each trio by twice the number of callable sites (to account for mutations transmitted from both parents; equation 1). We found the mean per-generation mutation rate in brown bears to be *μ_g_* = 0.84 × 10^-8^ per bp (95% CI: [0.69, 1.00]) for parents at an average age of 12.5 years across sexes. Table 1 shows the rate estimated for each trio separately. Our approach also produced consistent estimates of the mutation rate as the stringency of filters was increased (Figure S1), providing confidence in our results.

We did not find the mutation spectrum in the bear to be significantly different from the spectrum found in humans (χ^2^ test, *P* = 0.30; Figure 3). The transition-to-transversion ratio among mutations was 1.8 [1.3, 2.8], comparable to the expectation for SNPs in humans and other mammals. Similarly, we found that a substantial fraction of all mutations were CàT transitions at CpG sites (23%). We estimate the mutation rate at CpG sites in the brown bear to be 2.0 [1.3, 2.7] × 10^-7^ per bp per generation for parents at an average age of 12.5 years across sexes. This roughly order-of-magnitude higher mutation rate at CpG sites is consistent with previous estimates in other species.

**Figure 3.**
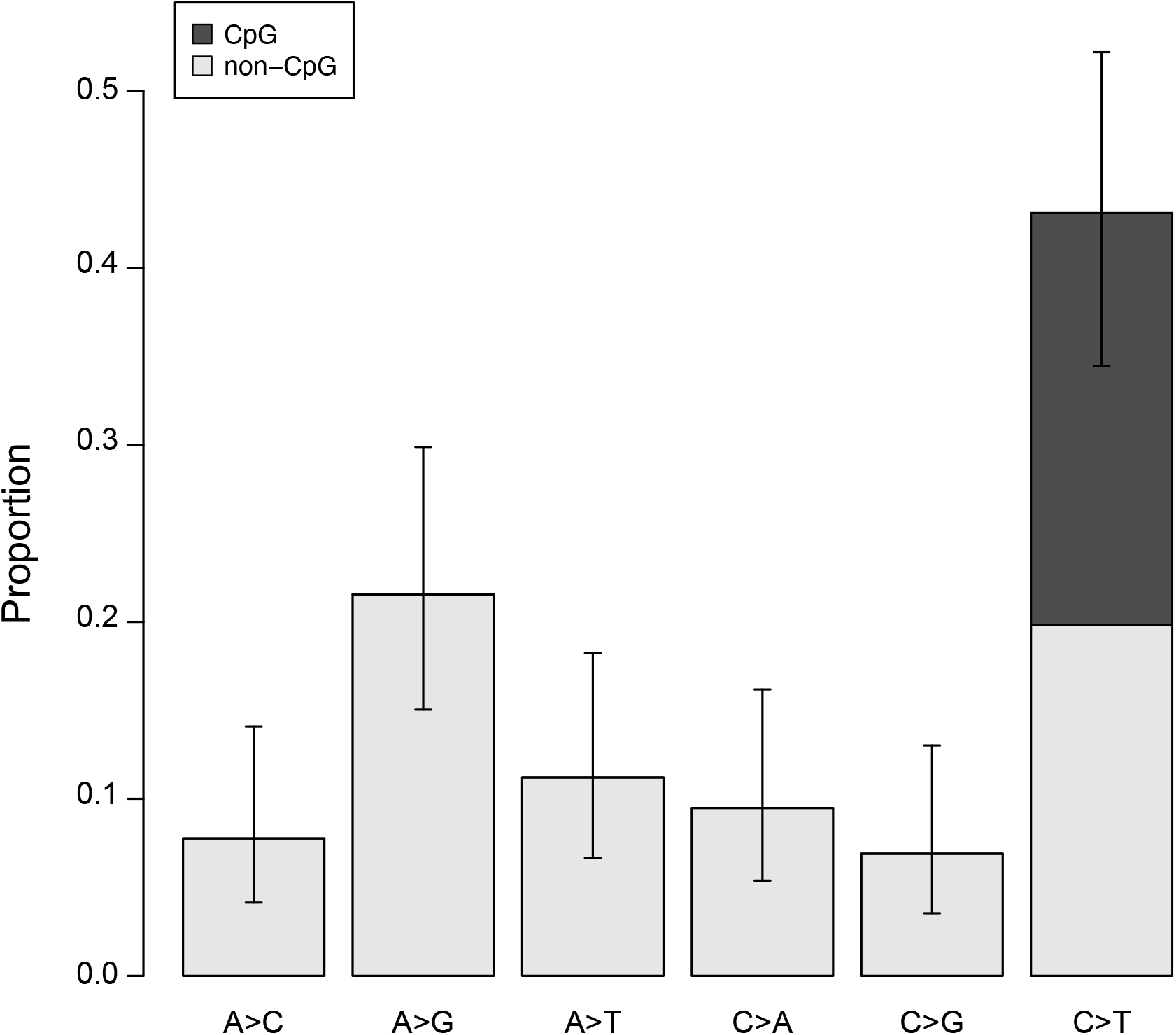
Bear mutation spectrum The proportion of each mutation class among bear trios, including their reverse complements. Dark gray region indicates the proportion of mutations occurring at CpG sites. Error bars show binomial 95% CI (Wilson score interval).

### Testing for an effect of hibernation on the per-generation mutation rate

The per-generation mutation rate for brown bears estimated here (*μ_g_* = 0.84 × 10^-8^ per bp) is lower than that observed in humans: 1.29 × 10^-8^ per bp for parents with an average age of 30.1 years (Jónsson et al. 2017). However, parents in the bear trios from this study are less than half the average human age, and the age at puberty in brown bears is also less than half of what it is in humans (5.5 years; White Jr. et al. 1998). A direct comparison of these rates therefore captures differences in reproductive life history between species rather than the potential effects of hibernation on mutation rates. In order to test for an effect of hibernation, we calculate the expected per-generation mutation rate under a model that considers mutation accumulation postpuberty in males (i.e., with no hibernation). We used the “reproductive longevity” model of Thomas et al. (2018) to calculate the expected per-generation mutation rate, assuming that male bears follow the same mutation trajectory as humans after reaching puberty at 5.5 years of age. Using equation 4 above, we calculate an expected *E*(*μ_g_*) = 0.85 × 10^-8^ per bp (Methods). This expected rate is not significantly different from, and only 1% greater than, the observed rate. We therefore conclude that hibernation in brown bears does not appear to produce a detectable effect on the per-generation mutation rate.

In addition to an effect on the overall per-generation rate, hibernation could reduce the proportion of paternally derived mutations. Such a reduction may be expected if spermatogenesis experiences a seasonal pause, as suggested by the absence of expressible sperm during hibernation and the presence of testicular regression after the breeding season. We investigated this potential effect of hibernation by examining the parent-of-origin across individual mutations. We were able to phase 26 of the 115 total mutations using read-pair tracing (Supplementary Table 1). Of the phased mutations, 22 were transmitted by a male parent and 4 were transmitted by a female parent. This proportion of male-biased mutations (84.6%) is highly consistent with the proportion found in humans (80.4%; Jónsson et al. 2017), and not significantly different from this proportion (χ^2^ test, *P*=0.8). The results again show no detectable effect of hibernation on the male mutation process, with a degree of male-bias as expected from uninterrupted germline replication post-puberty.

### Comparisons using the per-year mutation rate estimated from phylogenies

In order to look for possible effects of hibernation over a longer time period, we compared the number of substitutions in brown bears to the number in a sister lineage without hibernation (pandas). If hibernation has slowed the rate of mutation accumulation, we expect to observe fewer substitutions in the brown bear compared to the panda. Under the standard assumption that for neutral mutations the substitution rate equals the mutation rate (Kimura 1968), this comparison allows us to compare the mutation rate per-year between hibernating and non-hibernating sister lineages.

To study substitution rates, we used 4886 genic alignments among brown bear, panda, ferret, and dog (Methods). These two outgroups allowed us to compare the tip branch lengths specific to the brown bear and panda. We compared synonymous substitutions per site to minimize the effect of selection, finding the brown bear substitution rate to be 12.4% lower than that of the panda. The average length of the tip branch leading to brown bear (since its common ancestor with panda) was *d*s=0.0212 substitutions/site (S.E. 2.2×10^-4^) and the average length of the panda branch was *d*s=0.0242 (S.E. 2.2×10^-4^).

As was the case for our trio-based estimates of the per-generation mutation rate, a direct comparison of substitution rates does not immediately provide evidence for an effect of hibernation. The panda has both an earlier puberty time and a younger average age at conception in the wild than the brown bear (Peng et al. 2001; Aitken-Palmer 2010; Kersey et al. 2010), each of which can increase the rate of substitution per year relative to the brown bear (Laird et al. 1969; Wu and Li 1985; Thomas and Hahn 2014; Gao et al. 2016). To account for these differences, we calculate the expected difference in per-year mutation rates under a reproductive longevity model. Such a model can tell us whether an effect of hibernation needs to be invoked to explain the lower substitution rate in the branch leading to brown bears.

To predict the per-year mutation rate in both brown bear and panda, we used equation 7 above, applying appropriate ages for each species (Table 2) and using common mutational parameters estimated from humans. We calculated expected per-year mutation rates in these two lineages using a range of values for age at puberty and average age at conception in the wild (Table 2). From this range of life history estimates for the two species, we predict the brown bear per-year mutation rate should be 7.7%-14.4% lower than that of the panda, assuming the absence of any effect of hibernation. Our results from the phylogenetic analyses (12.4% reduction relative to panda) falls squarely within this range. We therefore conclude that hibernation has not led to a measurable difference in per-year mutation rates between the brown bear and the panda.

**Table 2.**

| Species | Paternal age at conception | Age of puberty in males | Maternal age at conception | Expected mutation rate per generation ($\times 10^{-8}$) | Expected mutation rate per year ($\times 10^{-9}$) |
| --- | --- | --- | --- | --- | --- |
| *Ursus arctos horribilis* | 13 | 5.5 | 6 | 0.85 | 0.89 |
|  | 13 | 4.5 | 6 | 0.89 | 0.93 |
|  | 11 | 5.5 | 6 | 0.77 | 0.91 |
|  | 11 | 4.5 | 6 | 0.81 | 0.96 |
| *Ailuropoda melanoleuca* | 7 | 4 | 6 | 0.68 | 1.04 |

## Discussion

By sequencing multiple independent trios (Figure 1), we identified 115 *de novo* nucleotide mutations in brown bears. These mutations allow us to make inferences about the per-generation mutation rate, the mutation spectrum (Figure 3), the degree of male-biased mutation, as well as the presence of multinucleotide mutations (multiple closely spaced mutations that occur in a single generation; Schrider et al. 2011). Our estimate of the per-generation rate, 0.84 × 10^-8^ per bp for parents at an average age of 12.5 years, joins a large and growing list of species for which this important evolutionary parameter has been measured (Chintalapati and Moorjani 2020; Yoder and Tiley 2021).

Our most striking result is the absence of an obvious effect of hibernation on the mutation rate, despite the apparent cessation of spermatogenesis and testicular regression associated with hibernation (Figure 2). Comparisons to non-hibernating species for the per-generation mutation rate, the per-year mutation rate, and the degree of male-bias reveal no significant differences. After accounting for the fact that a 12-year-old bear should only transmit approximately as many mutations as a 20-year-old human (because both are seven years post-puberty), we do not find a lower per-generation mutation rate in brown bears. Similar comparisons of the per-year rate in brown bear with the panda (a non-hibernating species) that take into account differences in life history between these two species also revealed no differences. While our model calculations have assumed that many of the underlying mutation parameters are the same among species (see Methods), we have previously found them to be conserved among multiple mammals (Thomas et al. 2018; Wang et al. 2020; Wang et al. 2021a).

There are several non-exclusive mechanisms that can explain the absence of a clear difference in mutation accumulation between hibernating and non-hibernating species. One obvious explanation is that an increasing mutation rate with age and a male bias in mutation are not driven by continuing mitosis during male spermatogenesis. As mentioned in the Introduction (also see de Manuel et al. 2022), there are several patterns from whole-genome sequencing studies that do not appear consistent with the classical role attributed to male germline replication. Although continued male germline replication is an obvious correlate of many of the coarse patterns of mutation accumulation, data from whole-genome sequencing has also uncovered multiple finegrained patterns that do not fit with this hypothesis. A simple model in which some aspect of mutation repair differs between males and females across most of their lifespan would fit the general trends equally well, and do much to explain several seemingly paradoxical patterns (Gao et al. 2019; de Manuel et al. 2022). Further investigation of underlying mutational mechanisms may help to add further detail to this newer model.

Despite the allure of new possible biological models, there are several ways to explain our data in brown bears that are consistent with classical hypotheses for the role of spermatogenesis in mutation accumulation. First, although there is a huge reduction in testis size, and no sperm could be sampled from hibernating bears, spermatogonial cells may be replicating throughout the year. Only a subset of spermatogonial cells is actively dividing even in full-sized testes (Plant 2010), and the absence of sampled sperm may be due to a halt in spermiogenesis rather than spermatogenesis. Under this model, despite all outward appearances, hibernation would have no appreciable effect on male germline replication. A second possibility is that spermatogenesis fully halts during the first part of hibernation, but then accelerates every spring during testicular recrudescence. This explanation would require a puberty-like process that occurs every year, ensuring that the male germline maintains the same total number of cell divisions per year, regardless of hibernation status. Such a hypothetical mechanism of accelerated replication could also explain the appearance of male-bias in mutation number just after puberty in humans (Gao et al. 2019), and the marked similarity in the number of mutations just after puberty across a number of species (Thomas et al. 2018; Wang et al. 2021a). Finally, while we have offered a few explanations for major patterns of mutation accumulation, male germline replication could play an important, but smaller, role than is currently believed.

Models of mutation accumulation are an essential part of understanding the evolution of mutation and the mutation rate. Per-generation and per-year mutation rates are key to many evolutionary inferences, from estimates of divergence times to explanations for the maintenance of sexual reproduction. Understanding the factors underlying changes in these rates is therefore necessary for researchers to form a comprehensive picture of many aspects of evolution. Interestingly, the results presented here are easily accommodated by current models of mutation. Even for models that were explicitly constructed with male replication in mind (e.g., Thomas and Hahn 2014; Amster and Sella 2016; Gao et al. 2016; Thomas et al. 2018), a simple re-parameterization that is agnostic to the causes of male bias will almost always result in the same outcome. For example, the rate of mutation accumulation in males post-puberty, *μ_yM1_*, need not depend on cell division rates for the predictions used here to hold (equation 5 above). Despite the success of such models, one outstanding question is how much our analyses will suffer if models remain phenomenological, and how much our science will improve if we fully incorporate molecular mechanism. There are clearly different processes of accumulation for different types of mutation—for instance, there is no parental age effect nor male-bias for structural mutations (Brandler et al. 2016; Belyeu et al. 2021; Thomas et al. 2021). One overall goal for the field will therefore be a general model of the cellular mechanisms that drive mutation rate evolution, a goal that will benefit from mutation rate studies in a wide variety of organisms and for a wide variety of different mutation types.

## Acknowledgements

We thank Julia Lowe and Teann Manser for assistance with analyses, and veterinarian Ahmed Tibary and the students and staff of the WSU Bear Center for bear handling and care.

## Data deposition

NCBI SRA for short reads SRR11336675-78, SRR11336682-85.

## Supplementary Figures

**Figure S1.**
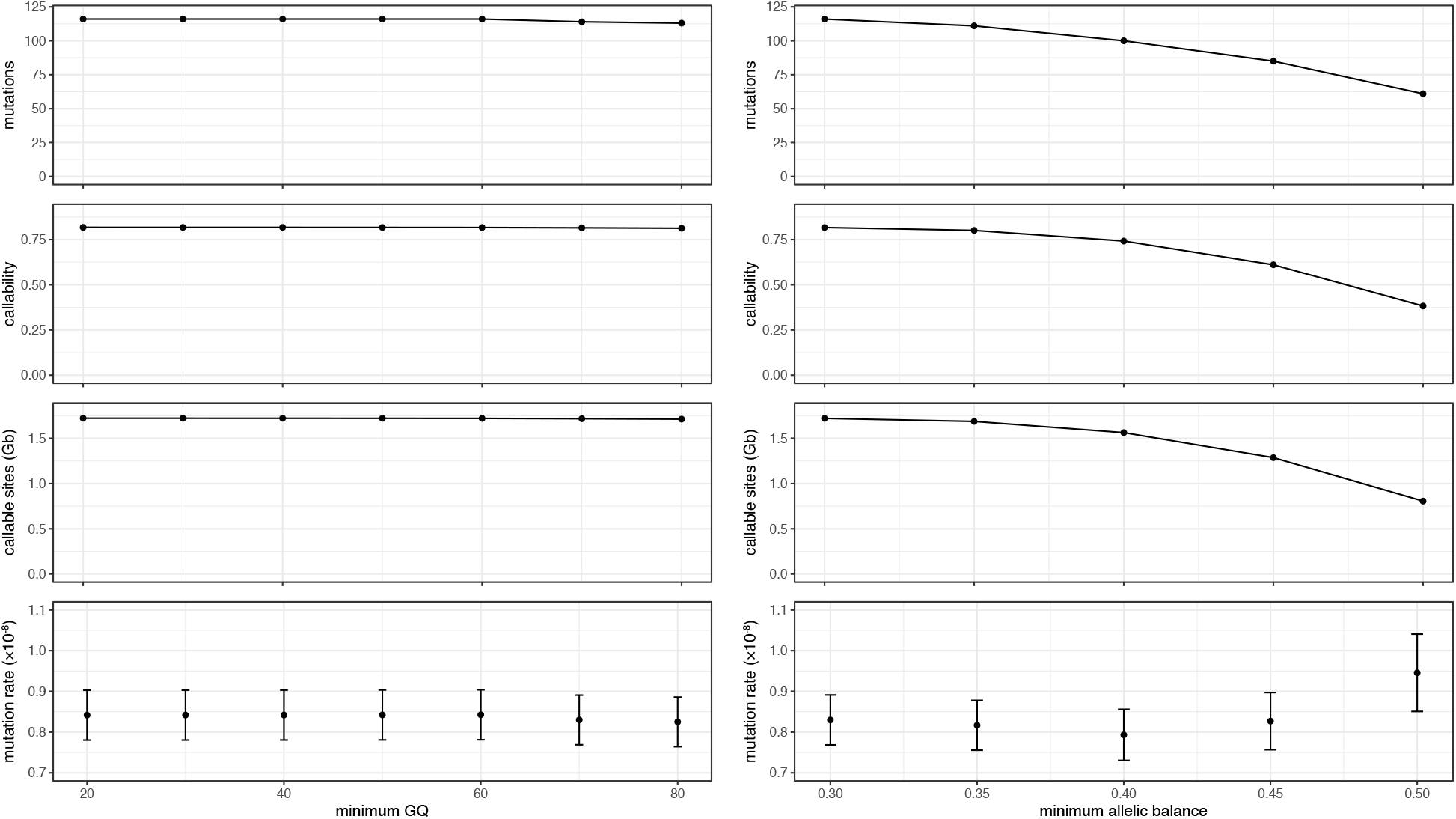
Mutation rates estimated with increasing filter stringency Mutation rates estimated with increasing stringency of minimum genotype quality (GQ) filtering, maintaining minimum allelic balance of 0.30 (left) and with increasing stringency of minimum allelic balance filtering, maintaining minimum GQ of 60 (right). Number of mutations (top), percent of the genome that is callable (callability), number of callable sites (in Gb) and final mutation rate are reported. Mutation rate error bars show 95% CI on estimated per generation rates under a Poisson model. Final mutation rates were calculated for sites with GQ > 60 and minimum allelic balance > 0.30.

**Figure S2.**
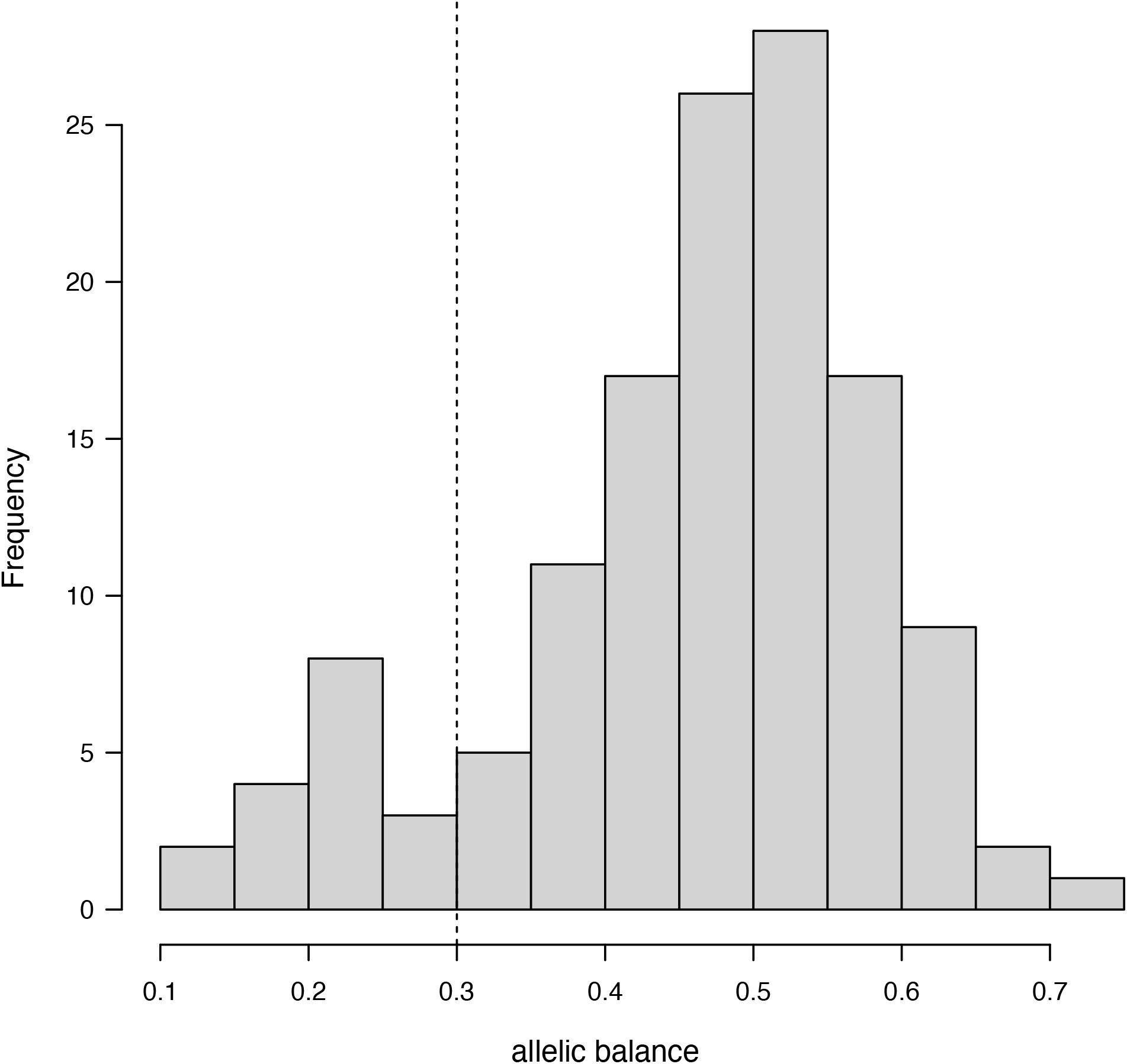
Distribution of allelic balance in candidate mutations. The observed allelic balance in candidate mutations that passed the first five filters (Methods). The sixth filter was that mutations had to have allelic balance > 0.3 (dashed vertical line). The expectation for true heterozygous sites is allelic balance = 0.5.

